# Microhomology-mediated CRISPR/Cas9-based method for genome editing in fission yeast

**DOI:** 10.1101/498402

**Authors:** Aki Hayashi, Katsunori Tanaka

**Affiliations:** Department of Bioscience, School of Science and Technology, Kwansei Gakuin University Sanda, Hyogo, Japan 669-1337

**Keywords:** CRISPR/Cas9, MMEJ, point mutation, knock-in, fission yeast

## Abstract

The CRISPR/Cas9 system enables the editing of genomes of numerous organisms through the induction of the double-strand breaks (DSB) at specific chromosomal targets. We improved the CRISPR/Cas9 system to ease the direct introduction of a point mutation or a tagging sequence into the chromosome by combining it with the microhomology mediated end joining (MMEJ)-based genome editing in fission yeast. We constructed convenient cloning vectors, which possessed a guide RNA (gRNA) expression module, or the humanized *Streptococcus pyogenes Cas9* gene that is expressed under the control of an inducible promoter to avoid the needless expression, or both a gRNA and Cas9 gene. Using this system, we attempted the MMEJ-mediated genome editing and found that the MMEJ-mediated method provides high-frequency genome editing at target loci without the need of a long donor DNA. Using short oligonucleotides, we successfully introduced point mutations into two target genes at high frequency. We also precisely integrated the sequences for epitope and GFP tagging using donor DNA possessing microhomology into the target loci, which enabled us to obtain cells expressing N-terminally tagged fusion proteins. This system could expedite genome editing in fission yeast, and could be applicable to other organisms.

## INTRODUCTION

The CRISPR/Cas9 system provides an outstanding prospect of manipulating genomes via DNA double strand break (DSB) induction and repair in numerous organisms from yeast to human (Dicarlo *et al.* 2013; Ran *et al.* 2013; Kondo and Ueda 2013). A guide RNA (gRNA)-Cas9 complex with high endonuclease activity is used to introduce the DSBs at designated points of a genomic region. The induced DSB is repaired by one of several repair pathways, mainly by non-homologous end joining (NHEJ), but also by homologous recombination (HR) and microhomology mediated end joining (MMEJ) (Ranjha *et al.* 2018). NHEJ often introduces unwilling insertions or deletions (indels), whereas the DSB is precisely repaired using HR. For genome editing, HR-mediated repair requires long homologous sequences at both ends of the DSB. In the budding yeast, *Saccharomyces cerevisiae,* 25–60 bp of the homologous sequences are adequate for chromosomal integration (Hayden and Byers 1992). In mammals, more than 200 bp of the homologous sequences are required (Rubniz and Subramani 1984). Therefore, preparation of long DNA fragments as donors for integration is necessary to perform precise genome modifications using the HR pathway. However, it is time-consuming or costly to prepare long donor DNA. To introduce point mutation or chromosomally tag in the gene of interest by the HR pathway, a series of synthesized oligonucleotides and sequential PCR steps are required to prepare the donor DNA.

The MMEJ pathway is an attractive pathway for genome editing using the CRISPR/Cas9 system since it uses short homologous sequences to repair the DSB. The MMEJ pathway is an alternative NHEJ pathway and it seems to be a type of backup repair pathway related to NHEJ and HR (Sfeir and Symington 2016). The NHEJ pathway requires the repair proteins including the Ku70–Ku80 heterodimer, XRCC4, DNA-dependent protein kinase catalytic subunit (DNA-PKcs), and DNA ligase IV. The MMEJ pathway, however, is independent on the Ku70–Ku80 heterodimer. In contrast to the HR pathway, the MMEJ pathway is an error-prone repair mechanism that requires only 5–25 bp of microhomology at the ends of the DSBs (Mcvey *et al.* 2017). Recent papers have reported MMEJ-mediated knock-in using TALEN and the CRISPR/Cas9 systems in worm, fish, and human cells with donor DNAs containing 6–40 bp of short homologous sequences at both ends (Nakade *et al.* 2014; Ata *et al.* 2018). It suggests that the MMEJ pathway can be adopted for effective genome editing in numerous organisms.

The fission yeast, *Schizosaccharomyces pombe,* is an extensively studied model organism since it shares similar chromosome structures and chromatin modifications with higher eukaryotes like human and mouse. By exploiting the relative ease of editing its genome, the fission yeast has offered many insights into critical cellular processes. Fission yeast is also a suitable model organisms for optimizing the conditions of genome editing mediated by the MMEJ pathway. Its merits, including rapid growth, single colony formation, and uniform genetic background, facilitate the determination of the optimal lengths of homologous sequences required for an MMEJ pathway in genome editing. It has been reported that MMEJ takes place at S/G2 phase and requires Exo1, Rad52, and Pol4 but not Ku proteins, suggesting that the MMEJ pathway in fission yeast is related with the HR pathway and single stranded annealing (SSA) pathway (Decottignies 2007).

The CRISPR/Cas9 genome editing system has been applied to fission yeast (Jacobs *et al.* 2014; Rodríguez-López *et al.* 2016; Fernandez and Berro 2016; Zhang *et al.* 2018). However, using the prevalent CRISPR/Cas9 system in fission yeast remains laborious particularly because of the complicated gRNA expression vector construction. The gRNA expression vector has inefficient restriction sites for inserting the gRNA target sequence, resulting in the difficulty of preparing the gRNA expression vector. Overexpression of Cas9 protein by a constitutive promoter inhibits the cell growth (Jacobs *et al.* 2014). In addition, preparing long donor DNA is labor-intensive.

To implement the genome editing-mediated MMEJ pathway, we improved the CRISPR/Cas9 system by generating more convenient vectors to express gRNA and the Cas9 protein in fission yeast. Using this system, we optimized the length of homologous sequences required for MMEJ-mediated genome editing. We found that 25 bp of the homologous sequences at both ends are sufficient for the introduction of point mutation and epitope tags into the genome using MMEJ-mediated genome editing. We also found that the double-stranded donor DNA could introduce the modification using MMEJ-mediated insertion at a high frequency. In addition, a one-step PCR product with microhomology at both ends could introduce a sequence at a target gene locus to generate the strain expressing N-terminally tagged fusion proteins. This protocol facilitates high-efficiency manipulation of the genome with less effort and allows for the investigation of genes in fission yeast and potentially in other organisms.

## MATERIALS AND METHODS

### Strains and media

The fission yeast strain PR109 (*h^−^, leu1-32 ura4-D18*) was used as the original strain for all transformations. All fission yeast media are described in Moreno *et al*. (1991). YE (low adenine medium) plate contains 0.5 % of yeast extract (Oxoid), 3 % of glucose and 2 % of agar. YES plate is a YE plate supplemented with 225 mg/L of amino acids and bases (leucine, histidine, adenine, uracil, and lysine (5S)). Edinburgh Minimal Medium + 5S (EMM5S) contains 11.77 g/L of EMM – Glucose (Sunrise Science Products), 2 % of glucose, and the supplements described above. Strains used in this study are listed in Table S1.

### Constructs

Plasmids and primers used for vector construction are listed in Tables S2 and S3. To construct a single gRNA expression vector, a 978 bp DNA fragment containing the partial *rrk1*^+^ promoter and a 966 bp DNA fragment of *rrk1*^+^ terminator were amplified from fission yeast genomic DNA with primers KT1910-KT1903 and KT1907-KT1913, respectively. A DNA fragment containing *rrk1*^+^ leader, gRNA scaffold, and the hammerhead ribozyme was amplified from pMZ283 (Addgene ID 52224) with primers KT1904-KT1906. The three fragments were mixed and amplified with primers KT1910-KT1913 to obtain 2,435 bp of concatenated DNA fragment. The fragment was cloned into PstI-SacI sites of pREP1 (Maundrell 1993) using the In-Fusion HD Cloning Kit (TaKaRa bio, catalog number 639649). The *BbsI* site of *LEU2* marker gene was disrupted by site-directed mutagenesis using primers KT1928-KT1929 with PfuUltra High Fidelity DNA polymerase (Agilent, catalog number 600385). To exchange *CspCI* sites with *BbsI* sites as gRNA target sequence cloning sites, inverse PCR was performed using primers KT1932-KT1933 with KOD-Plus-Ver.2 (TOYOBO life science, catalog number KOD-201) to replace a 36 bp sequence with a 47 bp sequence including two *BbsI* sites (pAH233). Humanized *Streptococcus pyogenes (Sp)*Cas9 was amplified from p*Sp*Cas9(BB)-2A-Puro (Addgene ID 62988) with primers KT1911-KT1912 and was cloned into the *XhoI-Bgl*II sites of pSLF273 (Forsburg and Sherman 1997) to construct a Cas9 expression vector that is inducible in the absence of thiamine (pAH235). To generate the *LEU2* marker vector expressing both the gRNA and Cas9 protein, the *nmt41p*-Cas9 fragment was amplified from pAH235 with primers KT2051-KT2052 and was cloned into the *Nco*I site of pAH233 using the In-Fusion HD Cloning Kit (pAH237). Site-directed mutagenesis was carried out to change *nmt41* promoter sequence of pAH237 to *nmt1* promoter using primers KT2087 and KT2088 (pAH236) and PrimeSTAR HS DNA polymerase (TaKaRa bio, catalog number R010A). To generate *ura4*^+^ marker vector expressing both the gRNA and Cas9 protein, the *Bbs*I site of *ura4*^+^ gene in pSLF173 and pSLF273 vector was disrupted with primers KT1975-KT1976 using PfuUltra High Fidelity DNA polymerase. The Cas9 DNA fragment amplified with primers KT1911-KT1912 was cloned into the *XhoI-Bgl*II sites of the plasmids. The *rrk1*^+^ promoter-gRNA scaffold was amplified from pAH233 with primers KT1973-KT1974 and was cloned into the *Mlu*I site of the Cas9 expression vector to construct pAH242 (*nmt1p*-Cas9 + gRNA) and pAH243 (*nmt41p*-Cas9 + gRNA).

### Construction of the gRNA expression vector

For the gRNA vector digestion by *Bbs*I or *Bpi*I (NEB, catalog number R3539S or ThermoFisher, catalog number ER1011), the gRNA vectors (pAH233, pAH237 and pAH243) amplified in *E. coli* were heated at 65 °C for 30 min and subjected to digestion by *Bbs*I at 37 °C for 1 h. After the digestion was validated through agarose gel electrophoresis, the enzyme was inactivated by the incubation for 20 min at 65 °C. The digested vectors containing the Cas9 gene were aliquoted in small volumes and stored at −80°C. The 20 bp of gRNA target sequences were determined by finding the PAM nearest to the mutation site. The number of off-target sites was searched using CRISPR direct (https://crispr.dbcls.jp).

To clone the gRNA target sequence into the gRNA scaffold, the guide oligonucleotides were synthesized as follows for ligation into the pair of *Bbs*I sites:

5’-caccNNNNNNNNNNNNNNNNNNN-3’

3’-NNNNNNNNNNNNNNNNNNNcaaa-5’

For annealing oligonucleotides, equal volumes of two oligonucleotides were mixed in a microtube (each 5 μl of 100 μM oligonucleotides and 5 μl of 10 × annealing buffer (100 mM Tris [pH 7.5], 500 mM NaCl, and 10 mM EDTA) in 50 μl reaction volume, incubated at 95 °C for 5 min and cooled down to 25 °C for 30 min in a thermal cycler. The ligation reaction was carried out at 16 °C for 30 min using ~20 ng of digested gRNA vector, 1/100 diluted annealed oligonucleotides (0.1 pmol), and 0.5 volume of the Ligation high Ver.2 (TOYOBO life science, catalog number LGK-201), followed by transformation of *E. coli DH5α.*

### Design of the donor DNA for introduction of point mutation and knock-in using the CRISPR/Cas9 system

Donor oligonucleotides for mutagenesis were designed on the basis of the cleavage site position digested by the gRNA/Cas9 complex. The mutation site is located at ±4 bp from the cleavage site. Both ends of 15, 20, and 25 bp sequences that contained mutation sites from the cleavage site were synthesized for carrying out the introduction of point mutations in the genome (KT1023-KT1024: 15 bp *swi6-W104A* donors, KT1959-KT1960: 20 bp *swi6-W104A* donors, KT2089-KT2090: 25 bp *swi6-W104A* donors, KT2168-KT2169: 25 bp *mrc1-S604A* donors). A 575 bp of PCR fragment containing *swi6-W104A* was amplified using the primers KT1957-KT1958 and the template pKT2150, a plasmid containing the *swi6-W104A* allele generated by site-directed mutagenesis. Over 100 ng of PCR product was used to make the *swi6-W104A* mutant using the HR pathway. For 2 × Flag insertion at the 5’ end of the *stn1*^+^ gene, KT2133 and KT2197 oligonucleotides were used as donors. For GFP knock-in at the 5’ end of the *reb1* gene, a 763 bp of the *reb1*-GFP fragment was amplified using the primers KT2127-KT2179 and the template pFA6a-GFP(S65T)-kanMX6 (Bähler *et al.* 1998).

### Transformation and genome editing

Transformations were performed using *S. pombe* Direct Transformation Kit (Wako pure Chemical, catalog number 296-64303) with some modifications, based on the lithium acetate/PEG method (Sabatinos and Forsburg 2010). For genome editing by two plasmids, sequential transformation was carried out. The host cells were transformed with pAH235 and plated on EMM5S – uracil-containing thiamine plates (5 μM thiamine), which represses Cas9 expression. The transformants were cultured with liquid EMM5S – uracil (without thiamine) medium at 30°C for 1 day to express the Cas9 protein. The overnight culture was then diluted in EMM5S – uracil medium, and the diluted cells were cultured until the numbers of exponentially grown cells reached approximately 1 × 10^7^ cells/mL at 30 °C. Mutagenesis and knock-in experiments were carried out by co-transformation. 1 × 10^6^ cells (10 μL) transformed with pAH235 were sequentially transformed using *S. pombe* Direct Transformation Kit with 0.5–1 μg of the gRNA plasmid, 2 μL of heat denatured 10 mg/mL salmon testes DNA, the oligonucleotides for genome editing (1 nmol for single-stranded oligonucleotides, 900 pmol of mixed single stranded oligonucleotides, and 900 pmol of double-stranded oligonucleotides), or ~125 ng of *swi6-W104A* PCR product (see Design of the donor DNA) for the HR pathway. For GFP knock-in at the 5’ end of the *reb1* gene, ~3 μg of PCR product was used for transformation. The cells spread on an EMM5S – leucine – uracil plate were incubated at 32 °C for 6–7 days. The small transformants were streaked on YES plates and incubated at 32 °C for 2 days to obtain single colonies (Rodríguez-López *et al.* 2016). To confirm the loss of plasmids from the cells, a single colony was streaked on YES, EMM5S–leucine and EMM5S – uracil plates. The cells that did not grow on each EMM selective medium plates were selected and stocked. For *swi6-W104A* mutagenesis, the colonies were streaked on YES plates containing of 10 μg/mL of thiabendazol (TBZ) to evaluate the TBZ sensitivity. Using the cells exhibiting TBZ sensitivity, colony PCR was performed to amplify the region of *swi6* gene with primers KT1957-KT1958 and the PCR products were subjected to sequencing. For *mrc1-S604A* mutagenesis and knock-in experiments, all procedures were performed similar to in *swi6* mutagenesis experiment, except for the confirmation of the mutant phenotype and the insertion confirmation by microscopic observation and electrophoresis. Since *mrc1* mutants exhibit hydroxyurea (HU) sensitivity (Tanaka and Russell 2001), the transformants were streaked on YES plates containing 5 mM HU to identify the *mrc1*-deficient cells introduced by the MMEJ-mediated genome editing. HU-sensitive cells were used as templates for colony PCR, and PCR products amplified with primers KT27-KT90 were subjected to sequencing. For knock-in at the 5’ end of the *stn1^+^gene,* colony PCR products amplified from the region near the first ATG with primers

KT2049-KT2050 were separated by 2 % agarose gel electrophoresis to check the Flag insertion. PCR products were sequenced to confirm the in-frame 2 × Flag insertion. For GFP knock-in at the 5’ end of the *reb1*^+^ gene, transformants were observed by fluorescent microscopy to see the GFP signals. Colony PCR products amplified from the region near the first ATG with primers KT2125-2126 were separated by 0.8 % agarose gel electrophoresis to check the insertion and subjected to sequencing.

For genome editing with a single vector expressing both a gRNA and the Cas9 protein, the cells cultured in EMM5S – thiamine medium were subjected to transformation. Transformation was carried out as described above. To obtain a homogenous mutant population, the transformants were streaked on EMM-selective medium plates (without thiamine) and incubated at 32 °C for 3 days to select the cells carrying the vector. The cells were streaked on YES plates at 32 °C for 2 days to get single colonies. To confirm the loss of vector, single colonies were streaked on YES and EMM-selective medium plates, followed by colony PCR and sequencing as described above.

### Immunoblot analysis

The protein extracts were prepared using alkaline-TCA method as previously described (Knop *et al.* 1999). 1 × 10^7^ cells growing in early log phase were harvested and used to prepare the cell extracts followed by Immunoblot analysis. The 3 × Flag tagged Cas9 protein and the 2 × Flag tagged Stn1 were detected using monoclonal anti-Flag M2 antibody conjugated with HRP (Sigma, A8592). Anti-Cdc2 (Santa Cruz, SASC53) was used as a loading control.

### Microscopy

The preparation of living cells expressing GFP fused proteins was carried out as described previously (Hayashi *et al.* 2009). The cells were cultured in EMM5S liquid medium at 25 °C, and mounted on a glass slide. The cover glass was sealed with silicon grease (Dow corning, HVG-50) to observe the living cell by fluorescence microscopy. A BX51 microscope and DP71 CCD camera (Olympus) were used for fluorescence image acquisition and analysis.

### Reagent availability

The plasmids generated in this study (pAH233, pAH235, pAH237, and pAH243) are available from Addgene (ID number 121436, 121437, 121438, and 121439) and National BioResource Project-Yeast (ID number FYP4232, FYP4233, FYP4234, and FYP4235, http://yeast.nig.ac.jp/yeast/top.xhtml).

## RESULTS

### The improved CRISPR/Cas9 system efficiently introduces mutations in fission yeast

To simplify genome editing using the CRISPR/Cas9 system in fission yeast, we modified the gRNA expression vector described in Jacob *et al.* (2014) (Figure 1A–D). The vector pMZ283, which has the *rrk1* promoter, a gRNA scaffold, and a hammerhead ribozyme, is effective for expressing gRNA in fission yeast. However, the *CspCI* cloning sites in the gRNA scaffold are digested inefficiently (Rodríguez-López *et al.* 2016), resulting in the difficulty of cloning gRNA target sequences. To solve this problem, we generated a vector that has *Bbs*I cloning sites for the gRNA target sequence (pAH233, Figure 1A). *Bbs*I sites have often been used to clone gRNA target sequences in other organisms, and two *Bbs*I sites avoid self-ligation owing to different cleavage sequences (Ran *et al.* 2013; Kondo and Ueda 2013). The pAH233 vector has two *Bbs*I sites in the gRNA scaffold instead of *CspCI* sites, which are sandwiched between 1.3 kb of the *rrk1* promoter/leader sequence and 1.0 kb of the *rrk1* terminator (Wilhelm *et al.* 2008). The annealed gRNA target oligonucleotides can be efficiently cloned at the *Bbs*I sites without phosphorylation.

**Figure 1.**
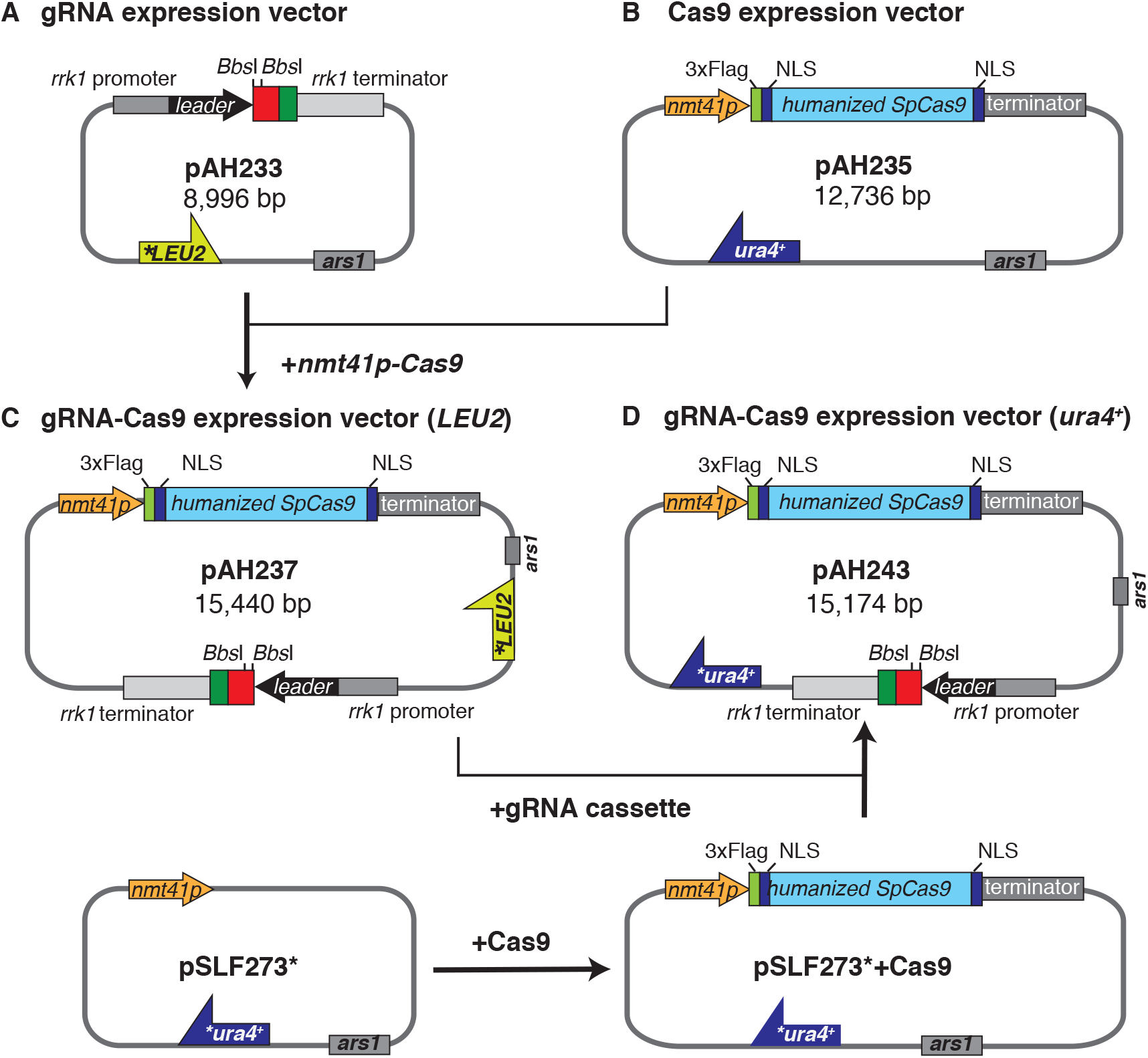
Construction of the gRNA and gRNA-Cas9 expression vectors. A. The gRNA expression vector (pAH233, *LEU2* marker). Red rectangle: gRNA expression cassette. Green rectangle: hammerhead ribozyme sequence. B. The *nmt41* promoter-driven Cas9 expression vector (pAH235, *ura4*^+^ marker). C. The gRNA-Cas9 expression vector (pAH237, *LEU2* marker). D. The gRNA-Cas9 expression vector (pAH243, *ura4*^+^ marker). Asterisks indicate silent mutations introduced by site-directed mutagenesis.

We also generated an inducible Cas9 expression vector. Because the expression of the Cas9 protein impairs the cell growth in fission yeast and the proper development of *Drosophila* (Jacobs *et al.* 2014; Port *et al.* 2014), it is critical to regulate the amount of Cas9 protein in the cells. The humanized *Sp*Cas9 protein fused to the 3 × Flag tag and two NLSs (Ran *et al.* 2013) was cloned into pSLF273 (*nmt41* promoter, *ura4*^+^ marker) vector to be expressed under the control of the *nmt41* promoter (Figure 1B, pAH235). We further generated a vector expressing both of a gRNA and the Cas9 protein to modify the genome through a single transformation using one marker gene, either *LEU2* or *ura4*^+^ (pAH237: *LEU2* marker, pAH243: *ura4*^+^ marker, Figure 1C and 1D). To ensure Cas9 expression, we generated the vectors that expressed the Cas9 using the *nmt41* promoter on each marker gene’s vector. Immunoblot analysis was performed using anti-Flag antibodies to validate the expression of the Cas9 protein in the cells carrying pAH233 in the presence or absence of thiamine (Figure 2A). The result showed that the expression of the Cas9 protein by the *nmt41* promoter was induced in the absence of thiamine, whereas its expression was repressed in the presence of thiamine (Figure 2A). We also analyzed the expression of the Cas9 protein from pAH237 and pAH243 in the absence of thiamine and observe that the Cas9 protein was stably expressed by the *nmt41* promoter (Figure 2B).

**Figure 2.**
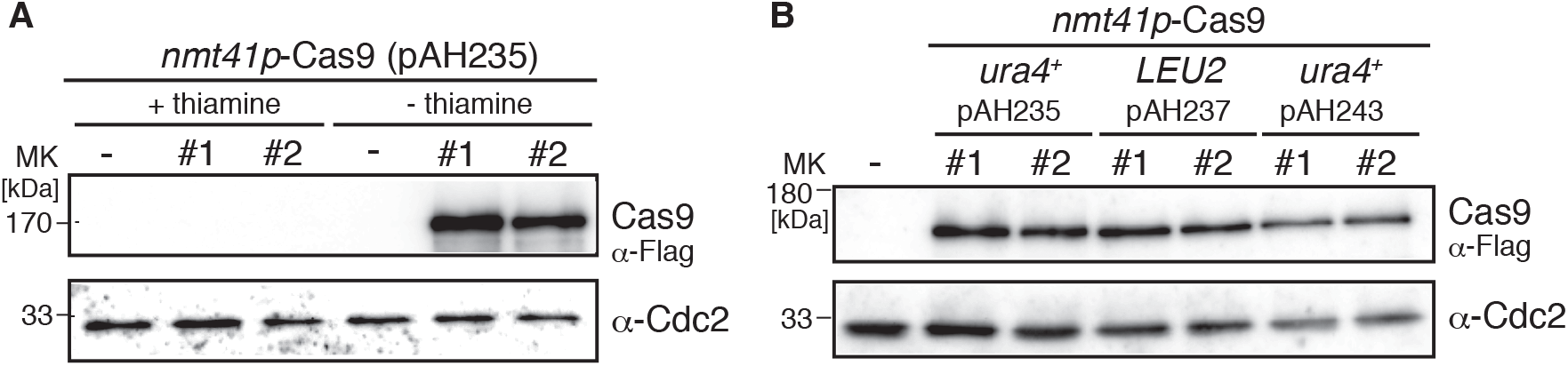
Immunoblot of the Cas9 protein expressed by the nmt41 promoter. A. The Cas9 protein was inducibly expressed by the *nmt41* promoter. Two independent transformants carrying pAH235 were cultured in EMM – uracil + thiamine or EMM – uracil – thiamine medium and were prepared as protein samples for SDS-PAGE (three lanes on the left; + 5 μM of thiamine (repressed condition), – thiamine condition (three lanes on the right). The extract of the whole cell carrying empty vector (pSLF273) was loaded as a negative control (–). Cas9 fused 3 × Flag protein was detected by α-Flag antibody. Cdc2 (α-PSTAIR) was used as a loading control. B. Detection of Cas9 proteins expressed by the *nmt41* promoter from the Cas9 and gRNA-Cas9 expression vectors. Two independent transformants carrying the indicated vectors were cultured in EMM-selective medium (without thiamine) to prepare the whole cell extracts followed by immunoblot analysis.

To examine the efficiency of the improved CRISPR/Cas9 system, we edit the *ade6*^+^ gene. The *ade6* gRNA target sequence was designed near *ade6-M210* mutation site to evaluate the mutation efficiency based on the frequency of the appearance of red color colonies (Jacobs *et al.* 2014). At the beginning, we tested a combination of *ade6*-gRNA vector (*ade6*-gRNA-LEU2, pAH244) and inducible Cas9 vector (*nmt41p-Cas9-ura4*^+^, pAH235). We sequentially transformed the wild type of *ade6*^+^ strain with the inducible Cas9 vector and then transformed it using the *ade6*-gRNA expression vector. Red colony color of the *ade6* mutant is because of the accumulation of red-colored intermediates in the adenine synthesis pathway. Obtained transformants were streaked on YE plates, a low adenine medium, to count the number of white and red/pink-colored colonies in order to confirm the introduction of an *ade6* mutation by the gRNA/Cas9 complex. The results revealed that 89 % of transformants expressed *ade6*-gRNA and the Cas9 protein showed red/pink color, whereas all of the cells carrying the control gRNA vector and the Cas9 vector showed white color (Table 1 and Figure S1A). We sequenced the *ade6* gene in eight red-colored transformants and found 1–3 bp indels in the *ade6* gene (Table S4), which indicates that the CRISPR/Cas9 system efficiently induced an *ade6* mutation comparable with that of a previously reported system (Jacobs *et al.* 2014). We also transformed wild-type cells expressing the Cas9 protein with the *rrk1* promoter-*Csp*CI-*ade6* gRNA cassette that has previously been used (pMZ284, derived from pMZ283, Addgene ID 52225) and cloned them into a *LEU2* marker vector (pAH234; pRE-pMZ284) to validate our modified system. The transformants carrying pRE-pMZ284 exhibited slow growth (Figure S2). It suggests that the cells had growth defects compared with the cells carrying pAH244, while the efficiency of *ade6* mutagenesis is 97 % (Table 1).

**Table 1.** Mutation frequency of *ade6*^+^ mutagenesis (two plasmids)

| vector | red colony# | white colony# | total | mutation freq. [%] |
| --- | --- | --- | --- | --- |
| no gRNA | 0 | 72 | 72 | 0 |
| *ade6* | 50 | 6 | 56 | 89 |
| pRE-pMZ284 | 32 | 1 | 33 | 97 |

Subsequently, we transformed the cells with the single vector expressing both the *ade6*-gRNA and the Cas9 protein. The wild-type cells were cultured in EMM5S without thiamine before the transformation. The cells were transformed with either the *LEU2*- or *ura4*-marked single vector and plated on EMM selective medium plates for 6–7 days. Transformants were streaked on YE plates to observe the colors of their colonies. The results revealed that the transformation with the *LEU2*-marked vector introduced the mutation at a high frequency (83 %, Table 2), whereas the transformation with the *ura4*-marked vector introduced the mutation at low frequency (13 %, Table 2). The transformants were then streaked on YE or EMM selective medium plates to determine whether they lost the vector. Approximately 50 % of transformants lost the vectors. The ratios of vector retention were following: *nmt41p-LEU2*: 11/33 (33 %), *nmt41p-ura4*^+^: 12/24 (50 %).

**Table 2.** Mutation frequency of *ade6*^+^ mutagenesis by single plasmid.

| Vector | red colony# | white colony# | total | mutation freq.[ % ] |
| --- | --- | --- | --- | --- |
| *ade6*-gRNA-*LEU2* | 19 | 4 | 23 | 83 |
| *ade6*-gRNA-*ura4<sup>+</sup>* | 5 | 35 | 40 | 13 |

We observed that transformants carrying one of the single vectors often exhibited a mosaic pattern of red and white cells (Figure S1B). To obtain homogenous cells, the transformants were streaked on EMM selective medium plates and incubated for 3 days to select the cells carrying the vectors. The cells carrying the vectors were then streaked on YE medium plates to obtain single colonies. The results revealed that all colonies exhibited the homogenous color (red color, Figure S1C). Comparing incubation on EMM-selective medium plate to incubation with no selective step, the frequency of the *ade6* mutagenesis was increased (*nmt41p-LEU2*: 7/8 [88 %], *nmt41p-ura4*^+^: 5/8 [63 %]). The result indicates that the cells that lose the gRNA/Cas9 vector promptly fail to express sufficient Cas9 protein to achieve genome editing. Therefore, it appears necessary for the introduction of the DSBs to regulate the duration of time required to express the gRNA/Cas9 complex.

### Introducing point mutations using a combination of the MMEJ pathway and the CRISPR/Cas9 system

The MMEJ-mediated knock-in has been reported in both TALEN and the CRISPR/Cas9 programmed systems in varied organisms, including frog, silkworm, and zebrafish as well as in mammalian tissue culture (Nakade *et al.* 2014; Ata *et al.* 2018), suggesting that the MMEJ pathway could be a useful pathway for genome editing in fission yeast. To validate the possibility and optimize the length of microhomologous sequences for genome editing, we attempted to introduce point mutations to appropriate regions of chromosome region through co-transformation of variable length of oligonucleotides and the gRNA-Cas9 vectors. We first designed a gRNA to target the *swi6*^+^ gene, which encodes an ortholog of heterochromatin protein 1. Swi6 binds to H3K9me via its chromodomain to form the heterochromatin (Nakayama *et al.* 2001; Jacobs and Khorasanizadeh 2002). Swi6-W104 is located in the chromodomain (Figure 3A), and Swi6-W104A mutant protein fails to interact with H3K9me, resulting in a silencing defect at *mat* locus (Hayashi *et al.* 2009). We transformed cells carrying the Cas9 expression vector with the *swi6* gRNA expression vector. Since *swi6Δ* mutant shows the sensitivity to TBZ (Keller *et al.* 2012), which is a microtubule-depolymerizing drug, the transformants were streaked on both a YES plate and a YES plate containing 15 μg/mL of TBZ to test whether the *swi6* gRNA/Cas9 complex could introduce the mutation in the *swi6* gene (Figure 3E).

**Figure 3.**
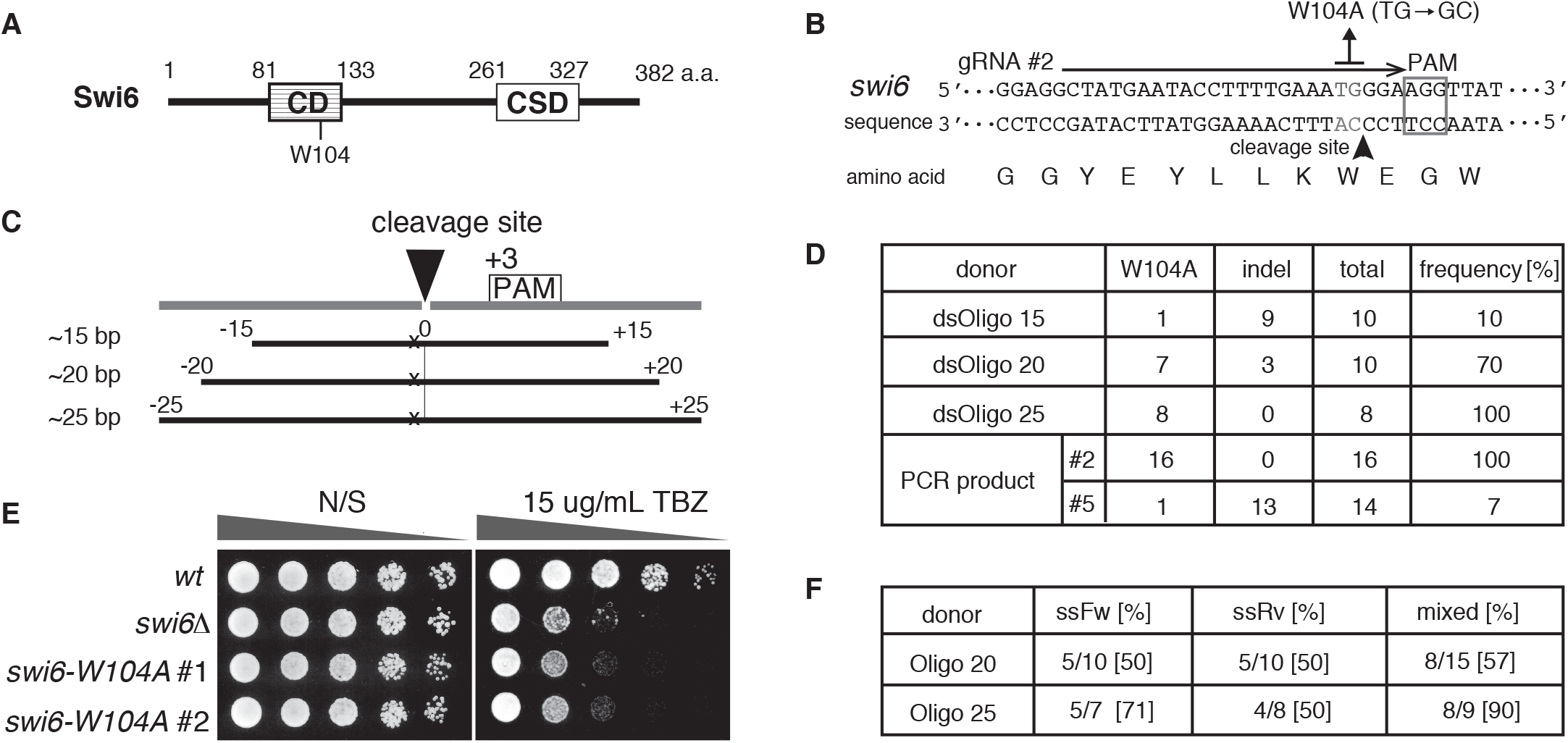
*swi6-W104A* chromodomain editing by the MMEJ-mediated genome editing. A. Diagram of Swi6 protein feature. CD: chromodomain, CSD: chromoshadow domain that is required for dimer formation. B. Sequences of the *swi6* gene and the gRNA target DNA for introducing the *swi6-W104A* mutation. The PAM sequence is flamed with a grey square, and the gRNA target sequence is indicated by a black arrow. The cleavage site is indicated by a black arrowhead. The grey letters (TG/AC) in the sequence indicate the nucleotides that will be substituted to GC/CG by editing (change the codon of tryptophan at 104^th^ to alanine). C. Design of the donor oligonucleotides for MMEJ-mediated mutagenesis. The mutation site (x) is located at +2 bp position from the cleavage site (closed triangle, at –3 bp position from PAM) digested by gRNA/Cas9 complex. The oligonucleotides are designed to contain the bidirectional 25 bp sequences from the cleavage site with the base substitution mutation (x) as indicated. D. The introduced frequency of the *swi6-W104A* mutation by the different lengths of dsOligos. The Cas9 expressing cells were co-transformed with both of the oligonucleotides and *swi6* gRNA #2 vector that possesses the target sequence near *swi6-W104.* The number of the cells described in total was analyzed to determine the mutation site by sequencing. As a control donor for the HR pathway, 575 bp length PCR product containing *swi6-W104A* mutation was co-transformed with the *swi6* gRNA #2 vector and also the *swi6* gRNA #5 vector that was designed to digest at +51 bp position from *swi6-W104.* E. The *swi6-W104A* mutant exhibits TBZ sensitivity. Five-fold-diluted cultures of the indicated strains were plated onto YES non-selective medium (N/S) and YES containing 15 μg/mL of TBZ and incubated for 2–3 days at 32 °C. F. The *swi6-W104A* mutagenesis by ssOligoFw, ssOligoRv, and mixed ssOligos. The number indicates the ratio of the number of the *swi6-W104A* mutants/total analyzed cells.

According to the results, 54 % of transformants exhibited TBZ sensitivity (14/26 transformants), indicating that the *swi6* gRNA/Cas9 complex had appropriately targeted endonuclease activity. To optimize the microhomology length for mutagenesis, we transformed the cells with both of *swi6* gRNA expression vectors and oligonucleotides containing different microhomologous length sequences near the *swi6-W104A* allele (Figure 3B and 3C). We designed the donor oligonucleotides that had ~15 bp, ~20 bp, or ~25 bp of sequences on either side of the cleavage site (represented as cleavage site 0 in Figure 3C, –3 bp from protospacer adjacent motif (PAM)). We co-transformed the Cas9 expressing cells with both of the annealed donor oligonucleotides and the *swi6*-gRNA vectors to introduce the *swi6-W104A* mutation. As a control, a 575 bp PCR fragment containing *swi6W-104A* allele was used as a donor for the mutagenesis using the HR pathway.

TBZ-sensitive colonies were used in direct colony PCR to amplify the *swi6-W104* region followed by sequencing to determine the mutation site. The results showed that 15 bp homologous oligonucleotides introduced a *swi6-W104A* mutation at 10 % frequency (1/10, Figure 3D and Table S4). However, most of the mutations introduced by the 15 bp homologous oligonucleotides were indel mutations, and some of mutants had rearrangements near the cleavage site (Table S4). Notably, 20 bp and 25 bp homologous oligonucleotides introduced *swi6-W104A* mutations at 70 % (7/10) and 100 % (8/8) of the frequency, respectively (Figure 3D). As expected, PCR fragments containing *swi6-W104A* mutation could introduce an appropriate mutation into the chromosome at 100 % (16/16) frequency. Therefore, even using very short oligonucleotides, the MMEJ pathway can introduce mutations at high frequency, which is comparable with the HR pathway. We also designed another *swi6* gRNA vector (+51 bp far from *swi6-W104* position, #5 in Figure 3D) and tested the frequency of introduction of *swi6-W104A* mutation by the HR pathway using same PCR products (Figure 3D). However, the mutation frequency was not high when the position of the DSB is relatively distant from the mutation site. Only 1 out of 14 transformants had the *swi6-W104A* mutation (7 %). The result highlights the significance of the positions of the DSBs digested by the gRNA/Cas9 complex in the introduction of high-frequency mutations.

Single-stranded oligonucleotides have also been reported to be effective in introducing mutations in other organisms (Dicarlo *et al.* 2013; Engstrom *et al.* 2009). We subsequently investigated the frequency of the mutagenesis by transformation of single-stranded oligonucleotides in fission yeast. We tested the ssOligoFw (sense strand), ssOligoRv (antisense strand), the mixed complemental ssOligos (ssOligoFw and ssOligoRv), and the annealed oligo (double-stranded Oligo (dsOligo)), which have 20 bp or 25 bp homologous sequences on either side of the cleavage site. The sequencing analysis revealed that both of ssOligoFw and ssOligoRv could introduce the *swi6-W104A* mutation (Figure 3F and Table S4). The ssOligoFw introduced the mutation at higher frequency than the ssOligoRv, implying a difference in repair pathways occurred within the sense and antisense strands. Remarkably, the mixed oligos and dsOligo could highly introduce the *swi6-W104A* mutation at 90 % and 100 % frequency (Figure 3F and Table S4).

To investigate the efficiency of the MMEJ-mediated mutagenesis at other loci of the chromosomes, *mrc1*^+^ gene was mutagenized by oligonucleotides using a similar procedure. The *mrc1*^+^ gene encodes mediator of replication checkpoint protein 1, which is required for Rad3-dependent activation of checkpoint kinase Cds1 in response to replication fork arrest (Tanaka and Russell 2001) (Figure 4A). Mrc1-S604 is located in SQ repeats, which are potential substrates of Rad3/Tel1 kinase, and the *mrc1-S604A* mutant exhibited HU sensitivity due to reduced Cds1-T11 phosphorylation, which is required for the interaction with Mrc1 (Xu *et al.* 2006). We generated an *mrc1* gRNA expression vector and transformed cells carrying the Cas9 vector with the gRNA expression vector and the oligonucleotides possessing 25 bp of homologous sequences adjacent to *mrc1-S604* including the mutation (Figure 4B). The transformants were streaked on YES plates containing 10 mM HU to estimate the frequency of the introduced mutation in *mrc1* gene (Figure 4D). The frequency of the transformants exhibiting HU sensitivity was very high (96 % of transformants (23/24) exhibited HU sensitivity). Sequence analysis was performed using colony PCR products to determine the mutation sites of the HU-sensitive transformants. The results showed that the frequency was as high as that for the *swi6* mutagenesis: 60 % for ssOligoFw, 20 % for ssOligoRv, 90 % for mixed ssOligos, and 100 % for dsOligo (Figure 4C and Table S4). The results suggest the effectiveness of the MMEJ-mediated genome editing method in fission yeast.

**Figure 4.**
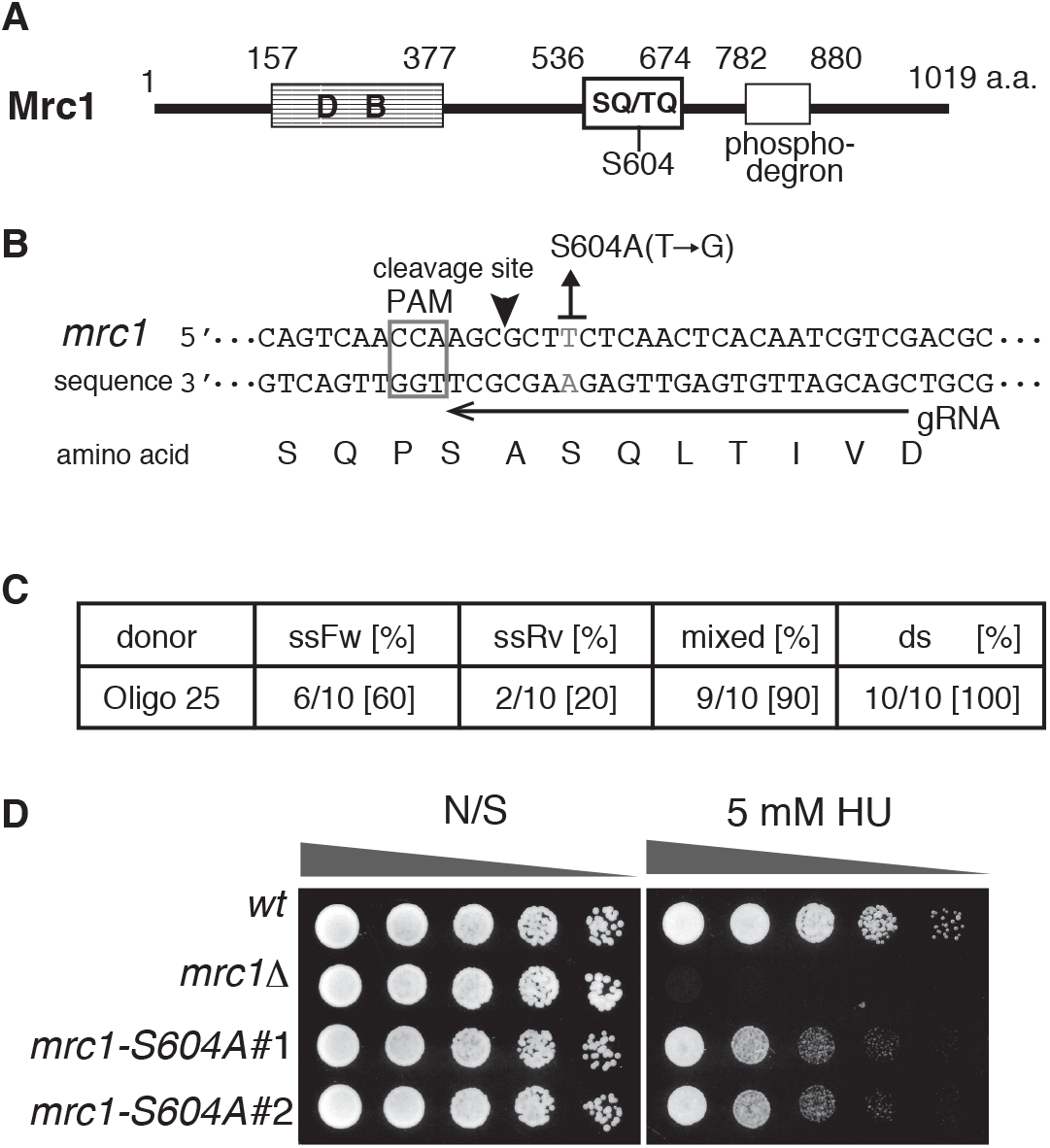
Introduction of the *mrc1-S604A* mutation by the MMEJ-mediated genome editing. A. Diagram of Mrc1 protein feature. DB: DNA binding domain, SQ/TQ: potential phosphorylation sites by Rad3/Tel1 kinase, phosphodegron: degradation domain by ubiquitin-proteasome. B. Sequences of the *mrc1* gene and the target gRNA sequence for introduction of the *mrc1-S604A* mutation. The PAM sequence is framed with a grey square, and the gRNA target sequence is indicated by a black arrow. The cleavage site is indicated by a black arrowhead. The grey letters (T/A) in the sequence indicate the nucleotides that will be substituted to G/C by editing (change the codon of serine at 604^th^ to alanine). C. The introduced frequency of the *mrc1-S604A* mutation by ssOligoFw, ssOligoRv, mixed ssOligos (mixed), and annealed ssOligos (ds). The number indicates the ratio of the *mrc1-S604A* mutants/total analyzed cells. D. The *mrc1-S604A* mutant exhibits HU sensitivity. Five-fold-diluted cultures of indicated strains were plated onto YES non-selective medium (N/S) and YES containing 5 mM of HU and incubated for 2–3 days at 32 °C.

### The MMEJ-mediated knock-in at the 5’ end of the gene combined the CRISPR/Cas9 system

In organisms such as silkworm and zebrafish, the MMEJ-mediated integration of fluorescent marker genes with TALEN and CRISPR/Cas9 system has been successful (Nakade *et al.* 2014; Ata *et al.* 2018). Therefore, we attempted to generate knock-in yeast cells using the MMEJ-mediated CRISPR/Cas9 system. Stn1 protein, one of the CST (Cdc13/Stn1/Ten1) complex proteins, is required for telomere protection and maintenance (Martin *et al.* 2007). We performed knock-in at the 5’ end of the *stn1*^+^ gene by transformation with oligonucleotides encoding a 2×Flag epitope tag sequence sandwiched between bidirectional 25 bp homologous sequences from the first ATG of the *stn1*^+^ gene (Figure 5A). The gRNA target sequence was selected from the sequence adjacent to the first ATG of the *stn1*^+^ gene, and the activity of the *stn1* gRNA/Cas9 complex was tested by transformation without the oligonucleotides. Sequence analysis revealed that 75 % of transformants had 1–3 bp deletion (6/8, Table S4), suggesting that the stn1-gRNA/Cas9 complex has high activity that is adequate for the introduction of the mutations in the *stn1*^+^ gene. Cells carrying the Cas9 expressing vector were co-transformed with both the *stn1* gRNA vector and donor oligonucleotides: ssOligoFw, ssOligoRv, mixed ssOligos, and annealed dsOligo. Electrophoresis analysis of colony PCR products shown in Figure 5B revealed that mixed ssOligos and annealed dsOligo effectively introduced the insertion (88 % and 96 % frequency, respectively, Figure 5C). In contrast, ssOligoFw and ssOligoRv introduced the insertion but at low frequency (21 % and 4 %, respectively, Figure 5C). Sequence analysis showed that the frequency of in-frame knock-in by transformation with mixed ssOligos and annealed dsOligo were high (82 % and 89 %, respectively, Figure 5C). Some of the knock-in strains contained the in-frame 3 × Flag insertion when the cells were transformed with the mixed ssOligos and dsOligo. Although the insertion frequency was low, ssOligoFw also introduced in-frame knock-in at high fidelity (80 % frequency, Figure 5C). We did not obtain the knock-in strain by the transformation with ssOligoRv (Figure 5C). We also found that the frequency of in-frame Flag tag insertion without indel was high (89 %) in the transformants obtained by transformation with dsOligo (Figure 5C and Table S4). The results suggest that the transformation with dsOligo would lead to precise knock-in without indel when using the MMEJ-mediated CRISPR/Cas9 system. We performed immunoblot analysis to confirm the expression of 2 × Flag-Stn1 protein. Flag epitope tag fused Stn1 protein was detected with anti-Flag antibody (Figure 5D), indicating that Flag epitope sequence was efficiently and precisely introduced at the 5’ end of the *stn1*^+^ gene.

**Figure 5.**
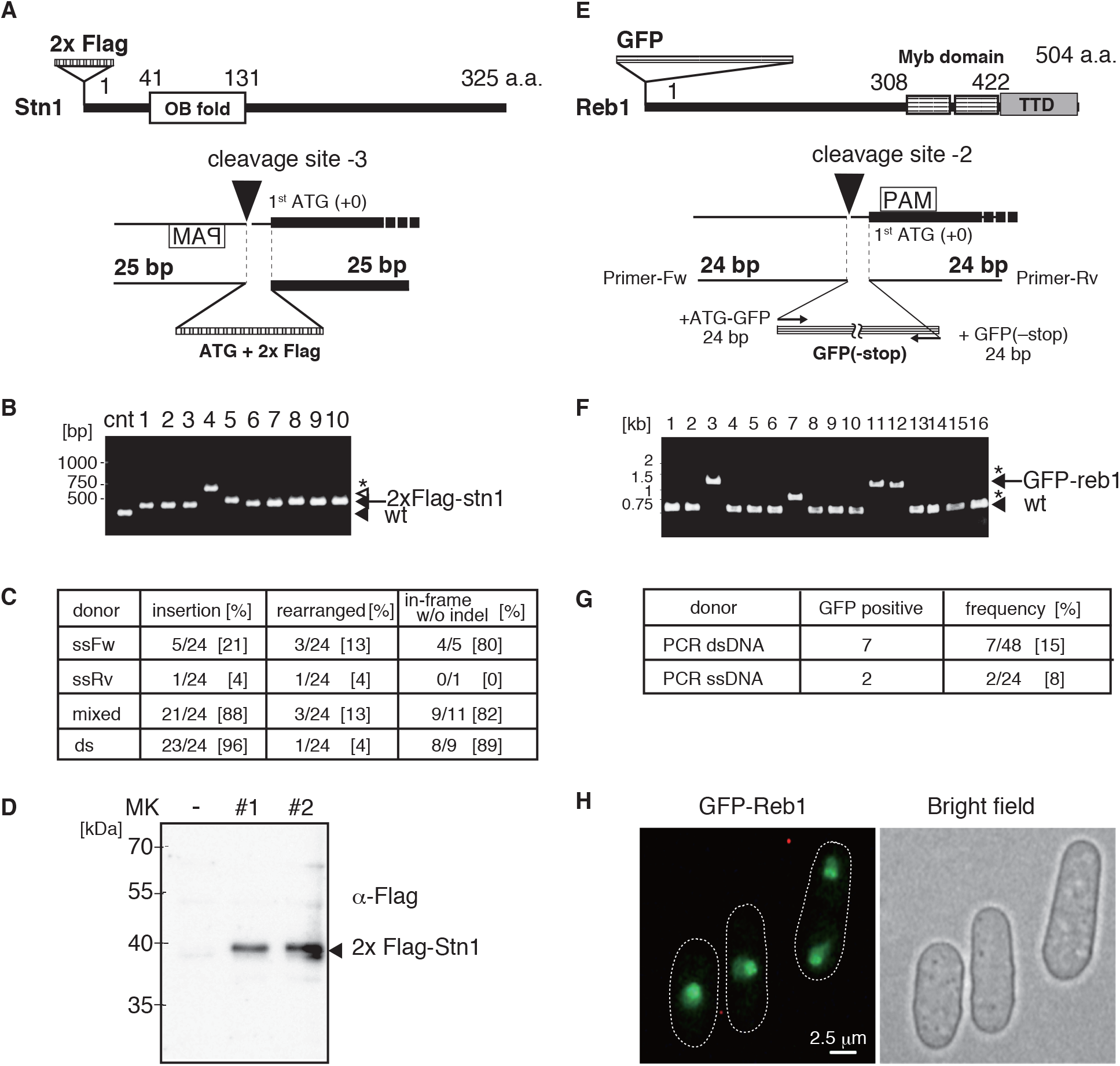
The MMEJ-mediated gene knock-in with CRISPR/Cas9 system. A. 2 × Flag knock-in at N terminal Stn1. Diagram of 2 × Flag tagged Stn1 protein and the design of oligonucleotides for the 5’ end of the *stn1*^+^ gene knock-in. The cleavage site is located at –3 bp position from first ATG. ATG + 2 × Flag repeat sequence is integrated by replacing 3 bp upstream sequence from first ATG. B. Electrophoresis of colony PCR products amplified the region at the 5’ end of the *stn1^+^.* Transformed cells with annealed oligos were used as templates. The 290 bp length of PCR product was used as a control (the bottom of black arrow, wt). The fragment containing 2 × Flag and 388 bp length of PCR product was detected (a black arrow at the top). White arrowhead indicates 3 × Flag inserted fragment. Asterisk indicates the fragment contained a part of the carrier DNA (salmon testes) inserted at the 5’ end of the *stn1^+^.* C. The frequency of 2 × Flag tagged *stn1*^+^ by single stranded oligonucleotides (ssFw, ssRv) and mixed ssFw and ssRv, and annealed ssFw and ssRv oligonucleotides (ds). Twenty-four independent transformants were analyzed to detect the insertion by electrophoresis. The ratio of rearrangement was indicated in the left column in the table. The denominator described at the column of in-frame without indel indicates the numbers of cells that were analyzed by sequencing. D. Immunoblot analysis of the 2 × Flag tagged Stn1 protein. Two independent 2 × Flag tagged strains were cultured to detect the expression of Stn1 protein with anti-Flag antibody. No tagged strain was loaded as a negative control (–). E. Diagram of GFP tagged Reb1 protein and the design of PCR primers for tagging. Fw primer contains 5’-UTR of *reb1*^+^ and the N-terminal GFP sequence. Rv primer contains the N terminal of *reb1*^+^ gene and C terminal GFP sequence without stop codon. The cleavage site is located at –2 bp from first ATG. F. Electrophoresis of colony PCR products at the 5’ end of *reb1^+^.* The 563 bp length of PCR product is used as a control (the bottom of black arrowhead, wt). A black arrow indicates the fragment containing GFP gene (1,326 bp). Asterisks indicate the fragments containing the rearranged sequence at the point of the 5’ end of *reb1*^+^ gene. G. The frequency of GFP fused Reb1 expressing cells obtained by genome editing. Heat denatured PCR product was used for transformation as a single-stranded DNA donor. H. The cellular localization of GFP fused Reb1 protein. Scale bar indicates 2.5 μm.

We further attempted to perform GFP gene knock-in at the 5’ end of the *reb1*^+^ gene using MMEJ-mediated knock-in. Reb1 is an rDNA-binding protein that is required for the termination of the transcription by RNA polymerase I (Zhao *et al.* 1997) (Figure 5E). Reb1 fused to GFP at the C terminus is located in the nucleus and the nucleolus (Hayashi *et al*. 2009). To generate a knock-in strain expressing GFP-fused Reb1 protein, we prepared a PCR fragment with GFP sandwiched between 24 bp upstream of the first ATG of the *reb1*^+^ gene and 24 bp downstream of the first ATG of the *reb1*^+^ gene (Figure 5E). Endonuclease activity of the *reb1* gRNA/Cas9 complex was evaluated on the basis of the frequency of the mutations introduced in the *reb1*^+^ gene. We performed a sequence analysis of colony PCR products amplified near the first ATG of the *reb1*^+^ gene and found that transformants had mutations in the *reb1* gene at a 75 % frequency (6/8, Table S4). The cells carrying the Cas9 vector were co-transformed with both of the *reb1* gRNA vector and PCR donor fragment, and transformants were analyzed as described above. The result showed that 7 out of 48 transformants had GFP gene insertion at a frequency of 15 % (Figure 5F and 5G). The heated denatured PCR fragment (single-stranded PCR product) could induce GFP insertion, but the frequency was lower than that of the double-stranded PCR fragment (2/24, 8 %, Figure 5G). Microscopic observation revealed that N-terminal GFP fused Reb1 was localized in the nucleus and the nucleolus (Figure 5H), as previously reported (Hayashi *et al.* 2009). The frequency of GFP insertion was lower than that of the Flag knock-in with synthesized oligonucleotides, suggesting that the length of donor DNA used in the integration influenced the insertion efficiency.

## DISCUSSION

Here, we describe the implementation of the CRISPR/Cas9 system in fission yeast to analyze the gene function with less-artificial genome modification. We improved the gRNA expression vector for ease of target sequence cloning. The regulation of the Cas9 expression levels using inducible promoter could introduce genome editing within a desired time and reduce a risk of the introduction of other mutations into the chromosomes by repressing Cas9 expression. It has been reported that Cas9 protein overexpression is toxic to cell growth (Jacobs *et al.* 2014). We also generated the vectors that expressed the Cas9 protein using the *nmt1* promoter, which could highly induce the transcripts more than the *nmt41* promoter could. However, the vectors introduced genome modification at low efficiency (15/23 (65 %): *nmt1p-LEU2,* 3/34 (9 %): *nmt1p-ura4*^+^, red colony/total transformants) and the retention of vectors in transformants was low (4/7 transformants (57 %): *nmt1p-LEU2,* 0/8 transformants (0 %): *nmt1p-ura4*^+^). Immunoblot analysis results demonstrated that the Cas9 protein expressed by the *nmt1* promoter was less stable and degraded in some cases (Figure S3). The *ura4*-marked vector particularly seemed highly eliminated from the transformants. Cells carrying the *nmt1* promoter vector would survive following the removal of the vector or halting of growth on selective medium. Some of the cells could be obtained by the integration of *ura4*^+^ gene at any loci of the chromosomes to prevent the expression of the Cas9 protein. It would cause the low frequency of genome editing with the vector possessing the *nmt1* promoter. We demonstrated here that the Cas9 protein expressed by the *nmt41* promoter is enough to introduce the mutation and the regulation of the duration of the gRNA/Cas9 complex expression is critical for the introduction of genome editing.

The MMEJ-mediated genome editing enabled us to prepare genome-modified cells with less time-consuming steps. There are established protocols in yeast for generating the gene deletion strains and the chromosomally tagged strains with the antibiotic-resistant markers using common vectors like pFA6a and its derivatives (Bähler *et al.* 1998). However, the insertion of the extra artificial sequences in some instances affects the authentic function of the gene of interest. Epitope tagging at the 3’ end of the essential genes sometimes fails because the tagged protein may lose or exhibit decreased activity.

For non-essential genes, tagged alleles may also demonstrate weaker activity compared with wild-type proteins. Such effects may be due to lower levels of expression or lower mRNA stability when the fused gene uses non-self UTR sequences derived from the tagging vector. The CRISPR/Cas9 system enables the modification of the genome without introducing marker genes facilitated by the high efficiency of DSB formation at the desired chromosomal locus. In the present report, we combined the CRISPR/Cas9 system and the MMEJ pathway to perform genome editing with short oligonucleotides. In numerous organisms, point mutations and chromosomal tagging of genes are often generated via the HR pathway with a long DNA donors. Preparation of a donor DNA with long (200–500 bp) regions of homology requires serial PCR steps and, therefore, is thus time-consuming. Using long (~100 bp) oligonucleotides for PCR can save the time required for preparation of a donor DNA. However, the cost of synthesizing long oligonucleotides is high, and the shorter homologous sequence lengths result in lower targeting efficiency. The MMEJ-mediated genome editing method reported here could save time and costs by using short synthesized oligonucleotides as a donor DNA. We demonstrated that the MMEJ pathway is highly effective for the introduction of point mutations into two genes using the CRISPR/Cas9 system with ~50 bp of short oligonucleotides.

As a donor DNA, double-stranded DNA seems improve the efficiency of the MMEJ-mediated genome editing. The annealed *swi6-W104A* oligonucleotides possessing 25 bp homologous sequences demonstrated higher frequencies during the introduction of the *swi6-W104A* mutation (100 %, Figure 3D) than the ssOligos possessing 25 bp homologous sequences (ssOligoFw: 71 %, ssOligoRv: 50 %, Figure 3F). The experiments that introduced the *mrc1-S604A* mutation showed that the dsOligo possessing 25 bp of homologous sequences edited the genome at a high frequency (100 %), in contrast, ssOligos introduced the *mrc1-S604A* mutation at a 60 % (ssOligoFw) and 20 % (ssOligoRv) frequency (Figure 4C). For 2× Flag knock-in at the 5’ end of the *stn1*^+^ gene, the dsOligo possessing 25 bp of homologous sequences at both ends was precisely inserted at a high frequency (89 %, Figure 5C), however, both ssOligos introduced an in-frame insertion at low frequency (ssOligoFw: 21 %, ssOligoRv: 0 %, Figure 5C). These results suggest that the MMEJ-mediated genome editing in fission yeast is related to the HR pathway, which takes place at G2 phase.

The frequency of GFP tagging, however, is lower (15 %, Figure 5G) than the introducing frequency of the 2 × Flag knock-in at the 5’ end of the *stn1* tagging. The length of GFP gene is about 700 bp, however, which is much longer than the homologous sequences required for annealing to the target genomic loci (24 bp each). Compared with GFP fragment, the 2 × Flag the length of the 2× Flag repeats is 54 bp, and the homologous sequences used for annealing were bidirectional 25 bp each. It highlights the significance of the annealing efficiency. Although the molecular mechanism of the MMEJ pathway requires further elucidation, the combination of the MMEJ and the CRISPR/Cas9 system in fission yeast could be carried out using both of the HR and the SSA pathways (Decottignies 2007).

In addition to the annealing efficiency, the amount of a donor DNA transformed into a cell may influence the efficiency of knock-in at the target locus of the chromosome. GFP-*reb1* PCR product is 763 bp, and we did not obtain any insertion when we used 100−200 μg of PCR product for transformation. As for the HR introduced *swi6-W104A* mutation, 65–130 μg of PCR product (575 bp) was adequate for obtaining the mutant (*swi6* gRNA #2 vector, Figure 3D). The observation could also be explained by the high annealing efficiency of donor DNA since *swi6-W104A* PCR product has over 250 bp homologous sequence at both ends.

We also observed that the ssOligos could induce the genome editing, and the forward oligonucleotides exhibited higher frequency of precise genome editing than the reverse oligonucleotides (Figures 3F and 4C). Notably, the mixed ssOligos could induce the genome editing at high frequencies comparable with dsOligo, although the efficiency of indel production was slightly higher (Figures 3F and 4C and Table S4). The differences in the insertion and indel frequencies occasioned by the opposite directions of ssOligos could be caused by the functions of different repair/replication proteins at the microhomologous region due to the different orientations of Cas9 and gRNA binding (Engstrom *et al.* 2009; Lemos *et al.* 2018). We cannot exclude the possibility that the orientation of the transcripts could be affected to the difference of the editing frequency. It remains unclear how variably the repair/replication proteins act at microhomologous sequences near cleavage sites. As for the repetitive sequence insertion, we obtained 3 × Flag tagged *stn1* strain by transformation with both of mixed ssOligo and dsOligo (Figure 5B and Table S4). The DNA replication machinery might generate an extra copy of Flag repeat sequence after the knock-in occurs in the chromosome.

In this paper, we demonstrate that the MMEJ-mediated genome editing combined with the CRISPR/Cas9 system is a powerful tool for generating point mutations and knock-in strains without a selective marker gene. It may also be possible to improve the current system using the Cas9 nickase to induce single-strand breaks, which may reduce off-target effects and introduce the genome modification at a position distant from the cleavage site (Ran *et al.* 2013; Satomura *et al.* 2017). The TALEN-based genome editing system has a potential choice (Gaj *et al.* 2014) and a combination of TALEN- and MMEJ-mediated genome editing could be a promising approach for modifying desired loci in any organism. We envision that this method would be a generally useful tool for genome editing and would facilitate the precise analysis of biological and molecular functions/mechanisms of genes in numerous organisms ranging from plants, animals, to humans in future studies.

## ACKNOWLEDGEMENTS

We thank Drs. Nick Rhind, Jun-ichi Nakayama and Da-Qiao Ding for critical reading of the manuscript. We also thank Dr. Sinya Takahata for advice of the CRISPR/Cas9 system. pMZ283 and pMZ284 were gifts from Dr. Mikel Zaratiegui. pSpCas9(BB)-2A-Puro (PX459) V2.0 was a gift from Feng Zhang. A.H. was supported by a Grant for Women researcher in Kwansei Gakuin University. T.K. was supported by JSPS KAKENHI Grant Number JP16K15094 and JP15H04630.

## SUPPLEMENTARY MATERIALS

Figures S1–S3 and Tables S1–S4 in the supplementary file.

